# Formation of *S. pombe* Erh1 homodimer mediates gametogenic gene silencing and meiosis progression

**DOI:** 10.1101/787739

**Authors:** Ditipriya Hazra, Vedrana Andrić, Benoit Palancade, Mathieu Rougemaille, Marc Graille

## Abstract

Timely and accurate expression of the genetic information relies on the integration of environmental cues and the activation of regulatory networks involving transcriptional and post-transcriptional mechanisms. In fission yeast, meiosis-specific transcripts are selectively targeted for degradation during mitosis by the EMC complex, composed of Erh1, the ortholog of human ERH, and the YTH family RNA-binding protein Mmi1. Here, we present the crystal structure of Erh1 and show that it assembles as a homodimer. Mutations of amino acid residues to disrupt Erh1 homodimer formation result in loss-of-function phenotypes, similar to *erh1∆* cells: expression of meiotic genes is derepressed in mitotic cells and meiosis progression is severely compromised. Interestingly, formation of Erh1 homodimer is dispensable for interaction with Mmi1, suggesting that only fully assembled EMC complexes consisting of two Mmi1 molecules bridged by an Erh1 dimer are functionally competent. We also show that Erh1 does not contribute to Mmi1-dependent down-regulation of the meiosis regulator Mei2, supporting the notion that Mmi1 performs additional functions beyond EMC. Overall, our results provide a structural basis for the assembly of the EMC complex and highlight its biological relevance in gametogenic gene silencing and meiosis progression.

## Introduction

Members of the ERH protein family are small proteins found in metazoan, invertebrates as well as plants. These proteins are strongly conserved (no amino acid changes between frog and human proteins and only one difference between human and zebrafish proteins), arguing for strict evolutionary constraints and for a highly important function. This gene was originally identified 25 years ago from a genetic screen as a mutant enhancing the truncated wing phenotypes of fruit flies lacking the rudimentary (r) gene ^1^, which encodes for the enzyme catalyzing the first three steps of the pyrimidine biosynthesis pathway. However, the biological function of ERH remains unclear despite its strong abundance in tumors compared to human normal cells, making it a very interesting candidate for functional characterization ^2^.

A bundle of evidences points towards a role of ERH in mRNA synthesis, maturation and nuclear export as this nuclear protein has been shown to interact with: (1) FCP1, the specific phosphatase for RNA PolII C-terminal domain ^3^; (2) the transcription factor SPT5 ^3^; (3) PDIP46/ SKAR (now named POLDIP3, for Polymerase delta-interacting protein 3), which localizes to nuclear speckles, regions enriched in pre-mRNA splicing factors ^4, 5^; (4) SNRPD3, a subunit of the Sm complex, which is involved in mRNA splicing. ERH also interacts with CIZ1, a zinc finger protein acting as a DNA replication factor and present in replication foci ^6, 7^. ERH is necessary for chromosome segregation during mitosis, probably through its role on CENP-E mRNA splicing ^8, 9^. Indeed, in the absence of ERH, the CENP-E mRNA, encoding a kinetochore protein, is incorrectly spliced and pre-mRNAs are rapidly eliminated by the nonsense-mediated mRNA decay pathway ^9^. Such link between splicing defects, cell cycle arrest and mitotic defects has already been observed for the depletion of other splicing factors ^10^. In *Xenopus*, ERH has been shown to act as a transcriptional repressor and to interact with DCoH/PCD (dimerization cofactor of HNF1/pterin-4a-carbinolamine dehydratase), a positive cofactor of the HNF1 homeobox transcription factor, by yeast two-hybrids ^11^. ERH protein is absent in *S. cerevisiae* yeast but expression of human ERH in budding yeast stimulates filamentous growth in low nitrogen media ^12^. Interestingly, this phenotype is reminiscent of the phenotype observed upon expression of the RBP7 subunit of the human RNA polymerase II in yeast ^13^, arguing again for a potential role of ERH proteins in the control of mRNA metabolism.

Closely related proteins, sharing around 30% sequence identity with human ERH, are also present in *Schizosaccharomyces* such as *S. pombe* and in few other fungi ^14^. Recent studies performed in *S. pombe* have enlightened the role of Erh1, the ortholog of human ERH. Initially, the *ERH1* gene was identified as a suppressor of *sme2*,*Δ* phenotype, *i.e.* a meiotic arrest due to the lack of inactivation of Mmi1 during meiosis ^15^. Mmi1 is a YTH-family protein ^16^, which selectively recognizes RNA hexanucleotide motifs (*e.g.* UNAAAC) present in meiotic transcripts and triggers their nuclear retention and elimination by the nuclear exosome during mitosis ^17^. Upon meiosis onset, Mmi1 is sequestered in a nuclear dot by the *sme2/mei* long noncoding RNA, which is assisted by the master regulator of meiosis Mei2 ^18^.

Recent works showed that Erh1 and Mmi1 form a 2:2 stoichiometric complex dubbed EMC (for Erh1-Mmi1 complex) whereby two Mmi1 peptides are physically bridged by an Erh1 homodimer ^17, 19, 20^. EMC localizes to scattered nuclear foci in vegetative cells and associates with two distinct complexes ^19, 21^. The first one known as MTREC (for Mtl1-Red1 core) is composed of the zinc-finger protein Red1, the Mtr4-like RNA helicase Mtl1 and Pir1/Iss10 among others subunits ^15, 19^. MTREC cooperates with Mmi1 to mediate degradation of meiotic mRNAs by recruiting the Rrp6 subunit of the nuclear exosome ^22^. The second complex known to interact with EMC is the CCR4-NOT complex but despite its known function as a mRNA deadenylase, it is not involved in Mmi1-dependent meiotic mRNA clearance ^19, 21, 23, 24, 25^. Instead, it is required for the integrity of heterochromatin and regulates the abundance of Mei2 protein during mitosis through the action of its Mot2/Not4 E3 ubiquitin ligase subunit 21. Interestingly, both MTREC (PAXT in human cells) and CCR4-NOT complexes are conserved in human cells, suggesting that ERH may also interact with these complexes in human cells. Furthermore, human ERH can partially rescue the sensitivity to sorbitol but neither SDS nor hydroxyurea of *S. pombe erh1*,*Δ* cells ^14^ indicating that a partially conserved function between Erh1 and human ERH proteins.

Here, we describe the crystal structure of *S. pombe* Erh1 protein and compare it to the structures of metazoan ERH proteins that have already been solved as well as to the structure of the *S. pombe* Erh1-Mmi1 complex that has been solved while this work was in progress ^20^. We observe that Erh1 organizes as a homodimer in which the two monomers contact each other via hydrophobic interactions, consistent with recent work ^20^. Structure-guided mutational analysis shows that formation of Erh1 homodimer is critical for cell growth at low temperatures and for its functions in meiotic mRNA degradation and meiosis progression. Interestingly, an Erh1 mutant (Erh1_I11R,L13R_) defective for dimerization still associates with Mmi1 *in vivo*, suggesting that Erh1 monomer is sufficient for interaction with Mmi1 while formation of Erh1 dimer is essential for EMC function. We also show that Erh1 does not contribute to the Mmi1-dependent down-regulation of Mei2 in mitotic cells, indicating that Mmi1 exerts functions beyond its partnership with Erh1. Overall, our results provide a structural basis for Erh1 dimerization and underscore the biological importance of Erh1 homodimer formation during both the mitotic and meiotic cell cycles.

## Results and discussion

### *S. pombe* Erh1 crystal structure

Initial polycrystals of Erh1 protein with a 6-branches star shape were obtained from an initial large screen of crystallization conditions in the following condition (0.8 M ammonium sulfate; 0.1 Na citrate pH 4). Thanks to the use of a micro-focus beamline, a complete dataset of moderate quality could be collected by shooting on a single branch of the star. Larger crystals could be obtained by increasing the drop volume, varying the ammonium sulfate concentration and the buffer. From one of these crystals, we could collect a dataset of better quality (see Table 1 for dataset statistics) from which we could determine Erh1 structure by molecular replacement using the structure of human ERH as initial model and refine it to 1.95Å resolution.

**Table 1:**
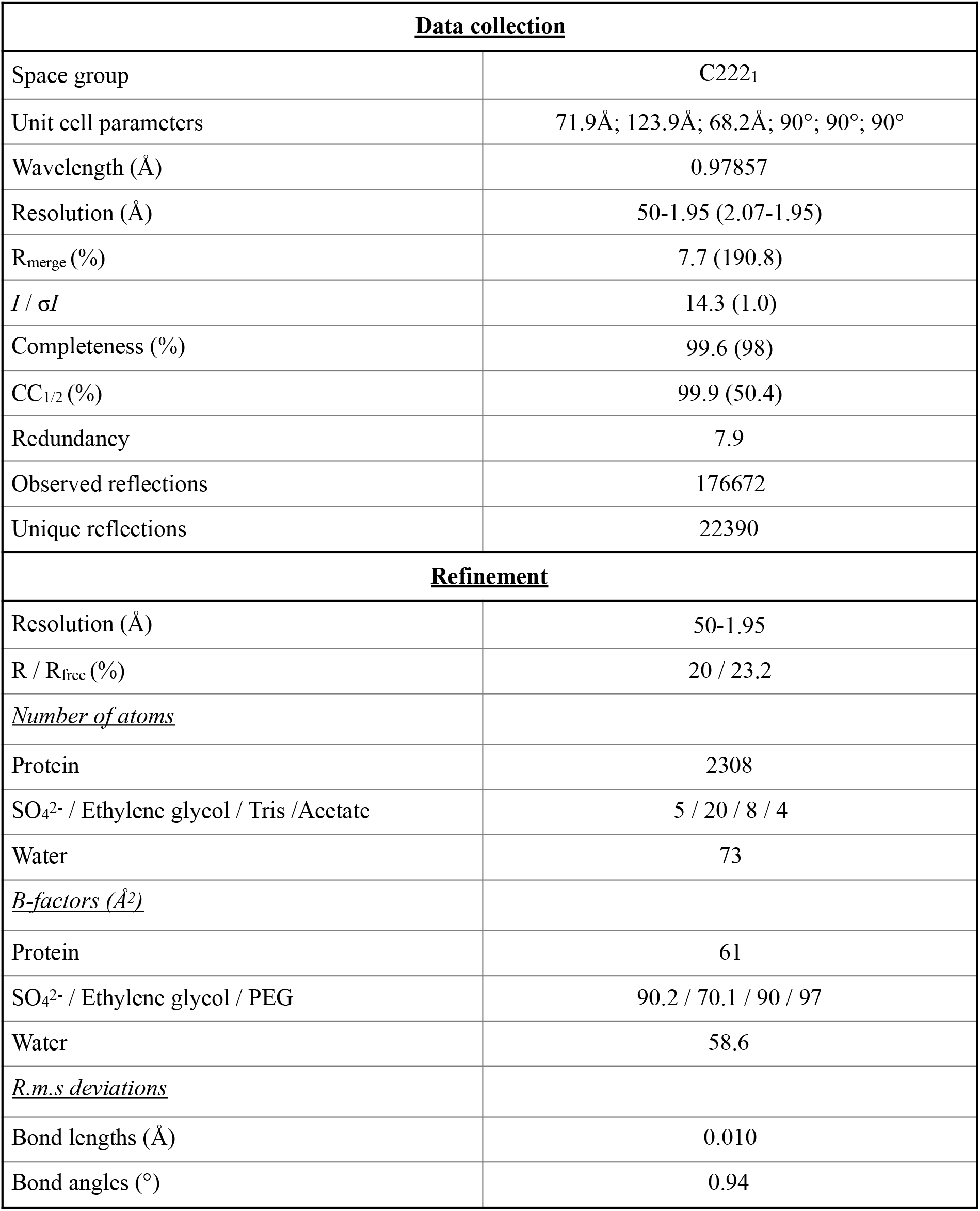
Data collection and refinement statistics

| **<u>Data collection</u>** |  |
| --- | --- |
| Space group | C222<sub>1</sub> |
| Unit cell parameters | 71.9Å; 123.9Å; 68.2Å; 90°; 90°; 90° |
| Wavelength (Å) | 0.97857 |
| Resolution (Å) | 50-1.95 (2.07-1.95) |
| R<sub>merge</sub> (%) | 7.7 (190.8) |
| *I* / $\sigma I$ | 14.3 (1.0) |
| Completeness (%) | 99.6 (98) |
| CC<sub>1/2</sub> (%) | 99.9 (50.4) |
| Redundancy | 7.9 |
| Observed reflections | 176672 |
| Unique reflections | 22390 |
| **<u>Refinement</u>** |  |
| Resolution (Å) | 50-1.95 |
| R / R<sub>free</sub> (%) | 20 / 23.2 |
| *<u>Number of atoms</u>* |  |
| Protein | 2308 |
| SO<sub>4</sub><sup>2-</sup> / Ethylene glycol / Tris / Acetate | 5 / 20 / 8 / 4 |
| Water | 73 |
| *<u>B-factors (Å<sup>2</sup>)</u>* |  |
| Protein | 61 |
| SO<sub>4</sub><sup>2-</sup> / Ethylene glycol / PEG | 90.2 / 70.1 / 90 / 97 |
| Water | 58.6 |
| *<u>R.m.s deviations</u>* |  |
| Bond lengths (Å) | 0.010 |
| Bond angles (°) | 0.94 |

Erh1 monomer folds as an α-β protein composed of a four stranded anti-parallel β-sheet and three α helices packed onto the same face of the β-sheet (Fig. 1A). This fold is similar to that of metazoan ERH (rmsd values of 1.1-1.9Å over 80 Cα atoms; ^12, 26, 27, 28^). The loop connecting helices α1 and α2 is not visible in our structure of Erh1. This most likely results from the intrinsic flexibility of this loop as shown for the corresponding region of metazoan ERH proteins by X-ray crystallography or NMR ^12, 26, 27, 28^. Three copies (protomers A, B and C) of Erh1 protein are present in the asymmetric unit. Protomers A and B are virtually identical (rmsd values of 0.27Å over 84 Cα atoms) while protomer C slightly differs as illustrated by its higher rmsd value when compared to the two other chains (1.3-1.4Å over 84 Cα atoms; Fig. S1A). The largest differences between protomer C and the two other Erh1 molecules present in the asymmetric unit occur at helix α2 (2.5Å translation along the helix longitudinal axis), at the hinge between helix α2 and the N-terminal extremity of strand β3, and to a lesser extent on helix α3 (1.3Å translation). It is noteworthy that human ERH adopts a conformation similar to that observed for Erh1 protomer C.

**Figure 1:**
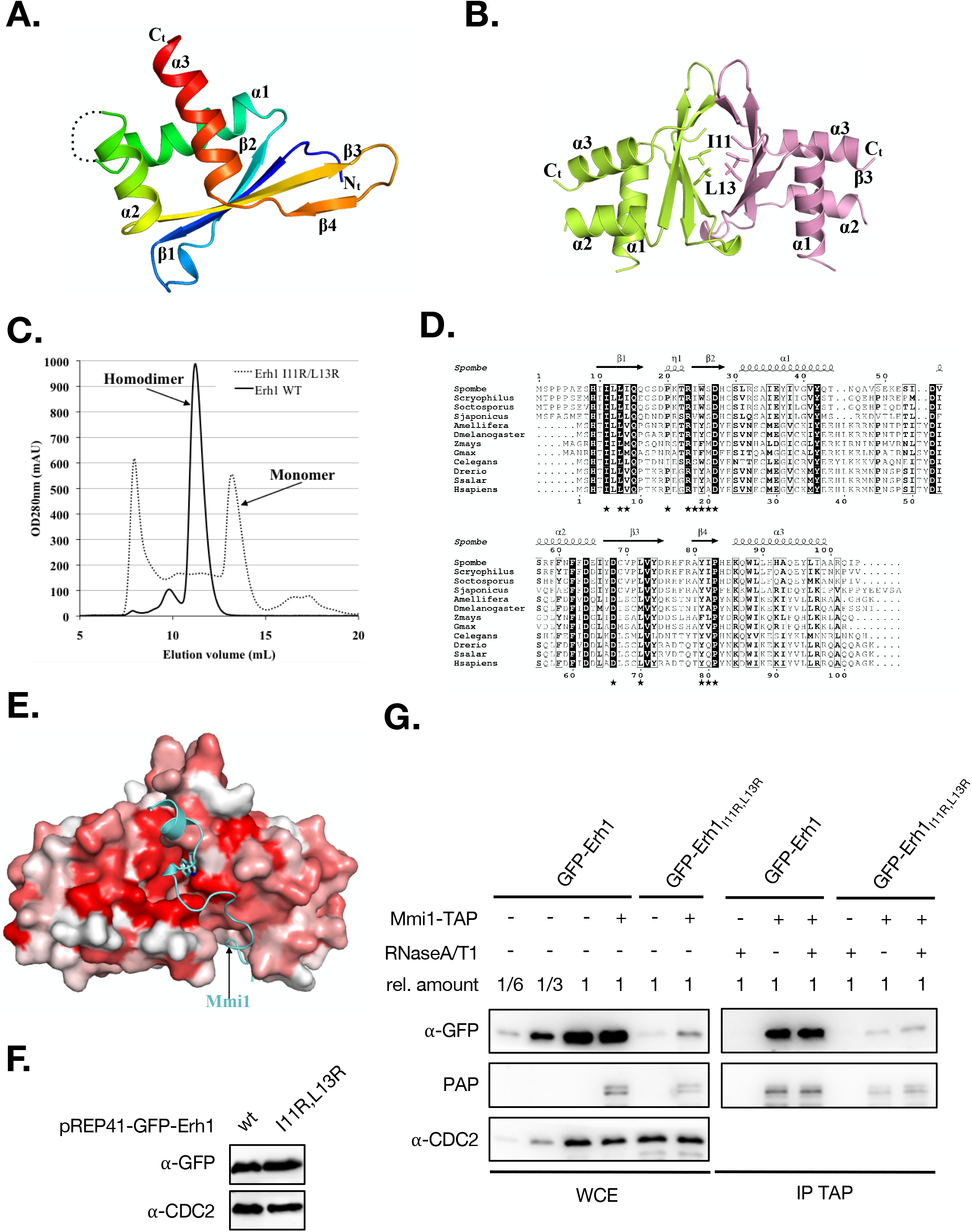
S. pombe Erh1 structure. A. Cartoon representation of Erh1 monomer. The protein is colored from blue (N-terminus) to red (C-terminus). The loop encompassing residues 47 to 54, which not defined in electron density maps, probably due to high flexibility, is depicted as a dashed line. B. Homodimer representation of Erh1. The Ile11 and Leu13 residues mutated in this study are shown as sticks. C. Analytical S75 size-exclusion chromatography profile of WT (solid line) and I11R/L13R double mutant (dashed line) Erh1 proteins. D. Sequence alignment of Erh1/ERH proteins from *Schizosaccharomyces* fungi (*S. pombe, S. cryophylus, S. octosporus* and *S. japonicus*), insects (*Apis mellifera* and *D. melanogaster*), plants (*Zea mays* and *Glycine max*), *C. elegans* worm and animals (*Danio rerio, Salmo salar* and *Homo sapiens*). Strictly conserved residues are in white on a black background. Partially conserved residues are boxed and in bold. This panel was generated using the ESPript server 37. Secondary structure elements detected from Erh1 apo structure are shown above the alignment. Residues involved in homodimer formation are indicated by black stars below the alignment. E. Sequence conservation score mapped at the surface of Erh1 homodimer. The conservation score has been calculated using the Consurf server ^38^ from the alignment shown in Fig. 1D. Coloring is from white (low conservation) to red (strictly conserved). The Mmi1 region interacting with Erh1 is shown as a cyan cartoon. The side chain for the Mmi1 tryptophan residue (W112) present at the heart of the interface is shown as sticks in panels A and B. F. Expression levels of dimeric GFP-Erh1 and monomeric GFP-Erh1_I11R,L13R_ in *erh1∆* cells. Western blot showing total GFP-Erh1 and GFP-Erh1_I11R,L13R_ levels expressed from the P_nmt41_ promoter (pREP41 vector) and obtained in denaturating conditions. An anti-CDC2 antibody was used for loading control. G. Monomeric Erh1_I11R,L13R_ protein still interacts with Mmi1. Western blot showing that both GFP-Erh1 and GFP-Erh1_I11R,L13R_ co-immunoprecipitate with Mmi1-TAP. (WCE) Whole Cell Extract; (IP) Immunoprecipitation.

### An evolutionary conserved homodimer

Among the three copies present in the asymmetric unit, protomers B and C associate to form a tight homodimer with a butterfly-like shape (Fig. 1B) while the protomer A forms a similar homodimer with a symmetry-related molecule (rmsd of 1.1Å over 160 Cα atoms). This dimeric state is consistent with the elution volume determined by size-exclusion chromatography (Fig. 1C). In the homodimer, the Erh1 β-sheet face from each monomer, which is not packed against α helices, interacts to form a β-barrel (Fig. 1B). Each monomer engages an area of 850 Å^2^ mostly formed by hydrophobic residues, which are strongly or strictly conserved within Erh1/ERH orthologues from fungi, plants, insects, worm and animals (Fig. 1D). This rationalizes that the Erh1 homodimer is reminiscent of those observed for metazoan ERH proteins (rmsd 1.4Å over 160 Cα atoms; ^12, 26, 28^).

An evolutionary highly conserved region present on the side of the homodimerization surface has been shown to participate in the interaction with Mmi1 (a short region encompassing residues 95-122; ^20^) while this work was in progress (Fig. 1E). Comparison between the Erh1-Mmi1 and our apo-Erh1 structures reveals that protomer C is strongly similar to the Erh1-bound structure and hence compatible with Mmi1 binding. On the contrary, significant differences support that protomers A and B are incompatible with Mmi1 binding (Fig. S1B-C). Most interestingly, the crystal structure of Erh1-Mmi1 complex reveals a 2:2 stoichiometry ^20^, confirming a previously proposed model of Erh1-mediated Mmi1 self-interaction ^17^.

As in this complex, each Mmi1-[95-122] peptide interacts with Erh1 on a region spread at the homodimer interface (Fig. S1B), we have decided to investigate the functional role of Erh1 homodimerization. We then simultaneously mutated two residues located at the homodimer interface (Ile11 and Leu13) into Arg to generate the Erh1_I11R,L13R_ double mutant with the aim of disrupting Erh1 homodimer. First, we have purified this mutant upon co-expression in *E. coli*. During purification, the Erh1_I11R,L13R_ mutant proved much less stable than the wild-type protein and had a tendency to aggregate as demonstrated by the presence of a large peak containing Erh1 and eluting in the void volume (7-8 mL) of a S75 size-exclusion chromatography (Fig. 1C). Furthermore, the 2 mL delay in the elution volume of a fraction of this Erh1_I11R,L13R_ double mutant (13.1 mL) compared to wild-type protein (11.1 mL) clearly indicates that the Erh1_I11R,L13R_ double mutant is monomeric in solution as anticipated (Fig. 1C).

To obtain additional insights into EMC complex assembly, we sought to determine whether Erh1 dimerization contributes to its association with Mmi1 *in vivo*. We generated *erh1∆* strains expressing GFP-tagged versions of the dimeric wild-type Erh1 or the monomeric Erh1_I11R,L13R_. Analysis of total protein levels under denaturating conditions indicated that the two forms of Erh1 were similarly expressed (Fig. 1F). When preparing native extracts for co-immunoprecipitation assays, however, we observed a significant decrease (roughly 6-fold less) in the total amount of Erh1_I11R,L13R_ when compared to wild-type Erh1 (Fig. 1G, WCE panel), likely as a consequence of decreased stability and in agreement with our observations during the purification of this mutant protein. Yet, surprisingly, Mmi1 still associated with Erh1_I11R,L13R_ (Fig. 1G, IP TAP panel), suggesting that Erh1 dimerization is not a prerequisite for its association with Mmi1, contrary to what could be assumed from the crystal structure of Erh1-Mmi1-[95-122] complex ^20^.

### Formation of Erh1 homodimer is required for gametogenic gene silencing

To investigate the role of Erh1 dimerization *in vivo*, we first analyzed its impact on *S. pombe* cell growth. Similar to *erh1∆* cells, the Erh1_I11R,L13R_ mutant displayed growth defects at all tested temperatures, especially 23°C, indicating that Erh1 dimerization is essential for its function (Fig. 2A).

**Figure 2:**
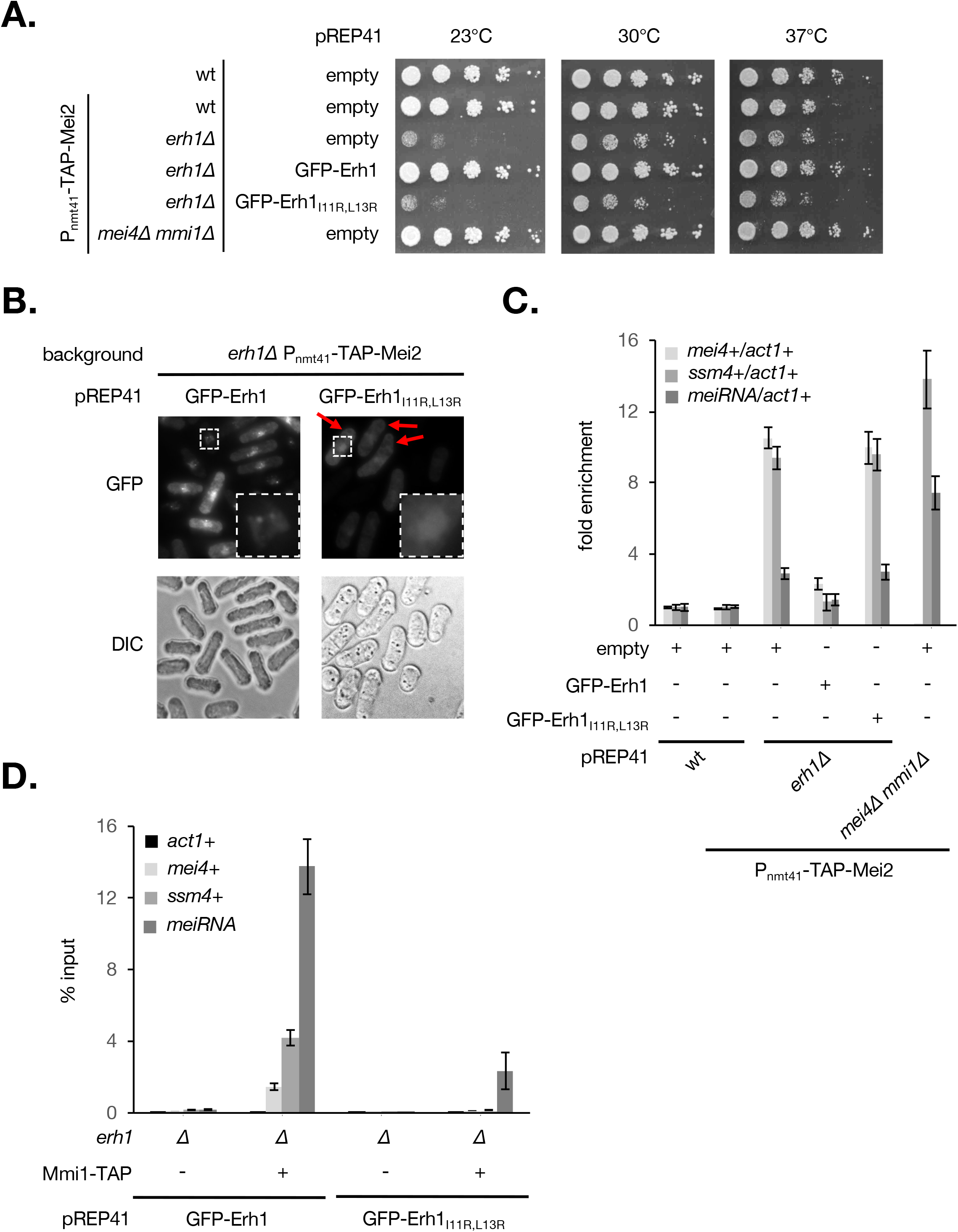
Erh1 dimerization is required for meiotic mRNAs recognition and degradation by Mmi1. A. The Erh1_I11R,L13R_ mutant phenocopies the deletion of *erh1*+ gene. Spotting assays at 23°C, 30°C and 37°C. Cells of the indicated genotypes were grown until mid-log phase and plated on EMM-LEU medium at an initial OD = 0.25 followed by 5-fold serial dilutions. B. Live cell microscopy of GFP-tagged Erh1 and Erh1_I11R,L13R_ in strains of the indicated genotypes. Cells were imaged by differential interference contrast (DIC) and with a GFP filter. Red arrows indicate Erh1_I11R,L13R_ cells for which GFP signals are similar to that of wild type Erh1. Squares with white dashed lines lying on the bottom right of GFP panels show enlarged images of the small squares. C. RT-qPCR analysis of the *mei4*+, *ssm4*+ and *meiRNA* transcripts in strains of the indicated genotypes. Signals were normalized to *act1*+ mRNA levels and expressed relative to wild-type cells. Error bars represent the standard deviation from four independent experiments. D.RNA-immunoprecipitation experiments in *erh1∆* cells expressing GFP-Erh1 or GFP-Erh1_I11R,L13R_. Shown are the enrichments (% input) of *act1*+, *mei4*+, *ssm4*+ and *meiRNA* transcripts upon pulldown of TAP-tagged Mmi1. Error bars represent the standard deviation of four independent immunoprecipitations from biological duplicates.

In vegetative cells, Erh1 and Mmi1 assemble in the EMC complex that localizes to nuclear foci ^17, 19, 20^. Live cell microscopy experiments revealed lower and diffuse GFP-Erh1_I11R,L13R_ signals when compared to GFP-Erh1 (Fig. 2B). Strikingly, nuclear dots were lost in the mutant, including in cells in which the GFP signal was similar to that of the wild type (Fig. 2B, red arrows). This supports the notion that Erh1 homodimer formation is a prerequisite for its confinement into nuclear bodies.

Erh1 cooperates with Mmi1 to target meiosis-specific transcripts for degradation by the nuclear exosome ^17, 19, 20^. To evaluate the role of Erh1 homodimer in this pathway, we measured the levels of Mmi1 RNA targets by RT-qPCR, including the *mei4*+ and *ssm4*+ meiotic mRNAs as well as the lncRNA meiRNA. Cells lacking Erh1 or expressing Erh1_I11R,L13R_ showed a strong accumulation of *mei4*+ and *ssm4*+ transcripts, while meiRNA levels were only partially increased when compared to the *mmi1∆* mutant (Fig. 2C). These results indicate that Erh1 dimerization is required for efficient Mmi1-dependent meiotic RNA degradation. To determine whether this was due to defective recruitment of Mmi1 to its targets, we analyzed the levels of meiotic RNAs co-precipitated with Mmi1. In cells expressing wild-type Erh1, Mmi1 efficiently bound to the *mei4*+, *ssm4*+ and meiRNA transcripts (Fig. 2D). Instead, in the Erh1_I11R,L13R_ mutant, the association of Mmi1 to meiotic mRNAs was abolished, while meiRNA still co-precipitated, albeit to a lower extent (Fig. 2D). Altogether, these data indicate that formation of Erh1 homodimer and hence proper EMC assembly is crucial for meiotic mRNAs recognition and degradation by Mmi1 but dispensable in the case of meiRNA. This different requirement for Erh1 dimer in the binding of Mmi1 to its RNA substrates might relate to the number of Mmi1 binding motifs (i.e. UNAAAC) within transcripts. Given *mei4*+ and *ssm4*+ mRNAs contain 8 and 7 binding sites, respectively, while meiRNA has up to 25 ^29^, it is possible that a pool of Mmi1 associates with the latter even in the absence of properly assembled EMC.

We previously showed that Mmi1 recruits the Ccr4-Not complex to promote ubiquitinylation and down-regulation of its own inhibitor Mei2, a master regulator of meiosis ^21^. To determine whether Erh1 also contributes to this regulatory circuit, we analyzed Mei2 levels in cells lacking Erh1 or expressing Erh1_I11R,L13R_. We found that, contrary to *mmi1∆* cells, Mei2 levels were not increased in mutants (Fig. S2), indicating that Erh1 does not contribute to Mei2 down-regulation in mitotic cells. This is also consistent with the notion that Mmi1 can exert functions independently of Erh1, as in case of transcription termination ^20, 30^.

### Formation of Erh1 homodimer contributes to meiosis progression

Erh1 not only suppresses the meiotic program in vegetative cells but also stimulates meiosis progression ^17, 19, 20^. To examine the requirement for Erh1 dimerization in meiosis, homothallic *erh1∆* cells expressing wild-type Erh1 or Erh1_I11R,L13R_ were exposed to iodine vapor, which stains the spore wall with dark color. Cells expressing wild-type Erh1 displayed strong staining intensity and high sporulation frequency, as indicated by the prevalence of asci (Fig. 3A). On the contrary, cells lacking Erh1 or expressing the Erh1_I11R,L13R_ mutant showed reduced intensity in staining, consistent with lower sporulation efficiency and asci formation (Fig. 3A). However, meiosis was not completely abolished as in *mei4∆ mmi1∆* cells, suggesting that the absence of Erh1 might be bypassed at least at low frequency.

**Figure 3:**
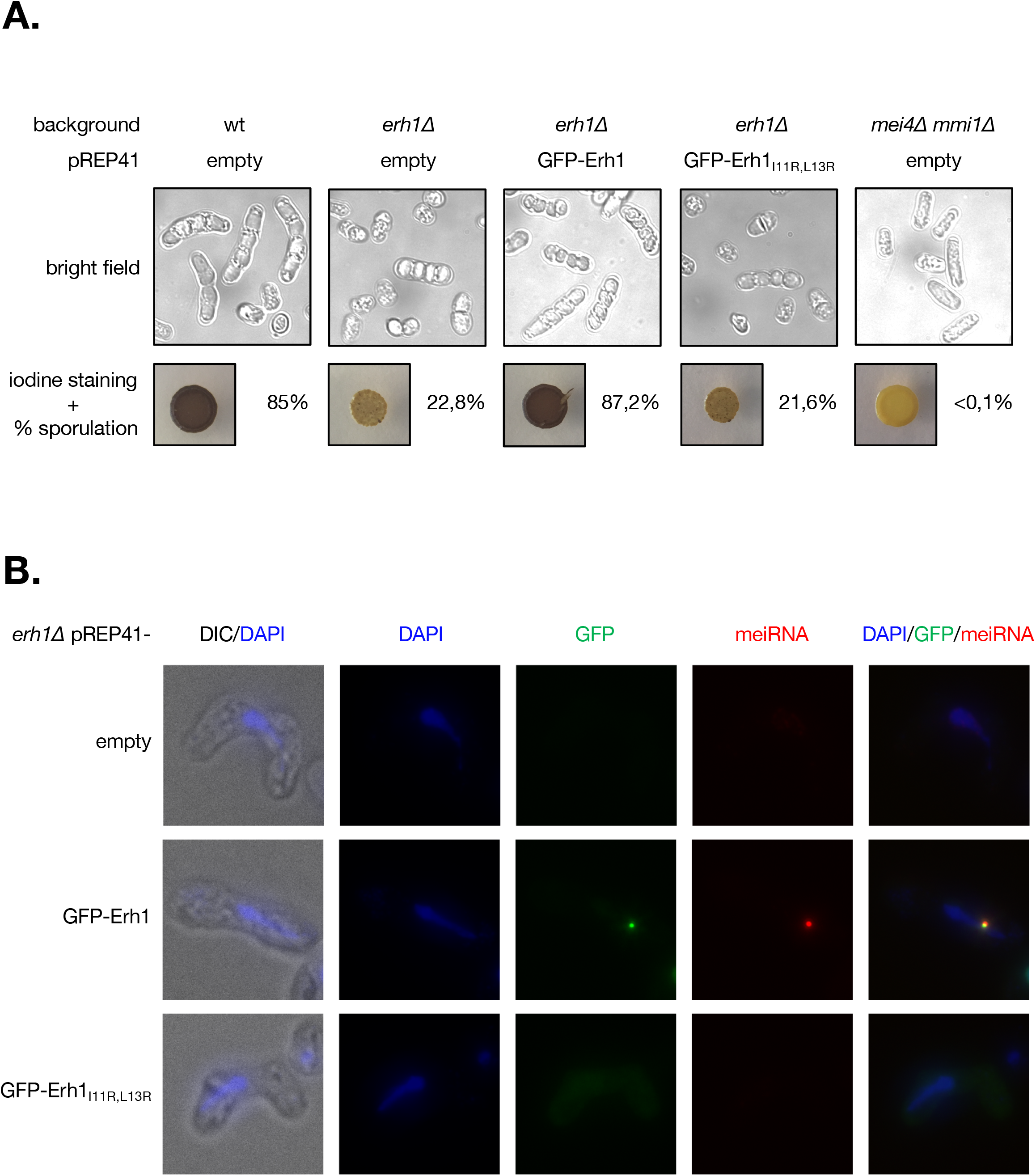
Erh1 dimerization is required for efficient meiosis progression and meiRNA dot formation. A. Homothallic strains of the indicated genotypes were spotted on EMM-LEU plates and incubated for 5 days at 30°C. The presence or absence of asci was determined by iodine staining and live cell imaging (bright field). The mating/sporulation efficiency is indicated for each strain and represents the percentage of asci among 500 cells. B. Representative images of meiRNA (red) detected by Single molecule RNA Fluorescence In-Situ Hybridization (SmFISH) in meiotic cells. DNA was stained with DAPI (blue). GFP-tagged wild-type Erh1 or Erh1_I11R,L13R_ were visualized in parallel. Images are shown as the maximum-intensity projections of Z-stacks.

Upon entry into meiosis, Mmi1 and Erh1 foci converge to a single nuclear dot associated with the meiosis regulator Mei2 and the lncRNA meiRNA ^18, 19, 31^. To determine whether Erh1 homodimer is necessary for dot formation, we probed meiRNA in meiotic cells by fluorescence *in situ* hybridization. Consistent with previous studies ^18, 19, 31^, meiRNA colocalized with wild-type GFP-tagged Erh1 in a unique nuclear dot (Fig. 3B). On the contrary, cells expressing GFP-Erh1_I11R,L13R_ failed to form both meiRNA and Erh1 dots (Fig. 3B). From these experiments, we propose that defects in Erh1 homodimer formation prevent the assembly of the EMC-meiRNA-Mei2 nuclear dots required for Mmi1 sequestration/inactivation during meiosis, thereby rationalizing impaired meiosis progression.

## Conclusion

In this study, we report the crystal structure of Erh1 and show that it assembles as a homodimer through hydrophobic interactions between N-terminal residues. Consistent with recent work 20, the structure also reveals a highly conserved region lying on the side of the dimerization domain and to which Mmi1 associates to form the heterotetrameric EMC complex. The strong similarity between Erh1 and human ERH homodimers (this study, ^20, 26, 28^) raises the possibility that the assembly of ERH-based multimeric complexes has been maintained throughout evolution for regulatory purposes, including modulation of gene expression. Whether Mmi1-related, YTH family RNA-binding proteins or other factors bind to ERH homodimers in metazoans to regulate cellular processes remains to be investigated.

Our functional analyses also highlight the biological relevance of Erh1 homodimer formation in gametogenic gene silencing. Importantly, Erh1 dimerization is dispensable for interaction with Mmi1 but essential for Mmi1 binding to meiotic transcripts, implying that RNA recognition by the YTH domain is not sufficient *per se* and that only fully assembled EMC complexes are functionally competent. This illustrates the cooperation between the C-terminal YTH domain and the N-terminal disordered region of Mmi1, to which Erh1 associates, for optimal binding to RNA substrates. The mechanistic basis for this is presently unclear but it is possible that the assembly of large macromolecular machines (*e.g.* EMC, MTREC, Ccr4-Not) facilitates protein-RNA interactions. In line with this, it is tempting to speculate that EMC and meiotic mRNA nuclear foci observed in vegetative cells ^17, 19^ may reflect multimerization driven by Erh1 dimers of such RNP complexes, gathering multiple transcripts in close proximity for efficient degradation.

Another important finding from our work is the requirement for Erh1 dimerization in Mmi1 sequestration/inactivation by the meiRNA-Mei2 dot during meiosis. The residual binding of Mmi1 to meiRNA observed in Erh1_I11R,L13R_ mitotic cells may not be sufficient for proper dot formation in meiosis. Therefore, Erh1 homodimer formation not only promotes Mmi1 function during vegetative growth but also contributes to its inhibition during meiosis. This cell-cycle dependent duality in functional outcomes may allow rapid changes in the activity of the complex without altering its expression levels or assembly. Whether this relates to the nature of the RNA substrate (meiotic mRNAs versus meiRNA) and/or additional factors (*e.g.* Mei2) remains to be determined. Regardless the precise mechanism, our study opens new perspectives to study the formation and activity of ERH homodimer-containing complexes in metazoan and to understand their relevance to human diseases ^10^.

## Materials and methods

### Cloning and protein expression

The gene encoding for Erh1 was amplified using *a S. pombe* cDNA library by PCR with oligonucleotides oMG511 and oMG512 (Table S1) using the Phusion High-Fidelity DNA Polymerase (Thermo) according to manufacturer’s instructions. The PCR products were further cloned into pGEX-6P1 vector using Fast Digest *BamH*I and *Xho*I restriction enzymes (Thermo Fisher Scientific) to generate the plasmid pMG921 encoding for a GST-tag fused to the N-terminal extremity of the full-length Erh1 by a 3C protease cleavage site (Table S1). The plasmid encoding for the Erh1_I11R,L13R_ double mutant was obtained by one-step site-directed mutagenesis of pMG921 using oligonucleotides oMG629/oMG630 (Table S1) to yield plasmid pMG945.

The Erh1 protein was expressed in *E. coli* BL21 (DE3) Codon+ cells upon transformation with pMG921 plasmid. Cultures were performed in 1 L of auto-inducible terrific broth media (ForMedium AIMTB0260) containing ampicillin (100 µg/mL) and chloramphenicol (25 µg/mL) first for 3 hours at 37°C and then overnight at 18°C. The cells were harvested by centrifugation at 4100 rcf for 45 minutes. The pellet was resuspended in 30 mL lysis buffer (20 mM Tris-HCl pH 7.5, 200 mM NaCl, 5 mM β-mercaptoethanol) in the presence of 100 µM PMSF.

Cell lysis was performed by sonication on ice, followed by lysate clearance by centrifugation at 20000 rcf for 45 minutes. The supernatants were applied on GSH-sepharose resin pre-equilibrated with lysis buffer. After extensive washing steps with lysis buffer and with a high salinity buffer (20 mM Tris-HCl pH 7.5, 2 M NaCl, 5 mM β-mercaptoethanol), the protein was eluted with elution buffer (20 mM Tris-HCl pH 7.5, 200 mM NaCl, 20 mM GSH, 5 mM β-mercaptoethanol). The eluted protein was next incubated overnight at 4°C with GST-3C protease under dialysis conditions in lysis buffer and then passed through GSH column to remove the GST-tag as well as the GST tagged 3C protease. The unbound proteins were subjected to size exclusion chromatography using HiLoad 16/60 Superdex 75 column (GE Healthcare Biosciences) pre-equilibrated with lysis buffer on an ÄKTA Purifier system (GE Healthcare Biosciences). The Erh1_I11R,L13R_ double mutant was purified using the same protocol.

### Crystallization, data collection and structure determination

Crystallization conditions were screened by the sitting-drop vapor diffusion method using JCSG+ screen (Molecular Dimensions) at 4°C by mixing 150 nL of concentrated protein (7.5mg/ ml) solution with an equal volume of reservoir solution in a 96-wells TTP plates (TTPlabtech). Initial hits corresponding to star shaped crystals were obtained in 0.8M ammonium sulfate; 0.1 Na citrate pH 4. For crystal optimization, hanging-drop method was used at 4°C, by mixing 1 µL of concentrated protein with 1 µL of reservoir solution. The best dataset was collected from crystals obtained in 0.3 M ammonium sulfate; 0.1 M Na acetate pH 3.8.

Prior to data collection, the crystals were quick-soaked in cryo-protectant solutions containing 15% (v/v) and then 30% ethylene glycol in corresponding well solutions and flash-frozen in liquid nitrogen. X-ray diffraction datasets were collected on both Proxima-1 and Proxima-2a beamlines at synchrotron SOLEIL (Saint-Aubin, France) and were processed with the XDS package ^32^. The dataset collected on Proxima-1 showed a higher resolution limit and was then used to determine and refine the structure of Erh1 protein (Table 1). The structure was solved by molecular replacement searching for 3 molecules in the asymmetric unit with the program PHASER ^33^. The initial model for molecular replacement was generated by the PHYRE2 server ^34^ using the crystal structure of human ERH (30% sequence identify) as template ^28^. In this model, the loop corresponding to residues 44 to 55 was removed as it is known to be highly flexible from the comparison of human and fruit fly ERH structures ^12, 26, 28^.

The final model was obtained by iterative cycles of building and refinement using COOT ^35^ and BUSTER ^36^, respectively (for final statistics, see Table 1). This model encompasses residues 6-46 and 55-100 for protomer A, 1-46 and 55-98 for protomer B (as well as 4 residues from the N-terminal tag) and 1-2, 7-47 and 54-98 for protomer C as well as 5 ethylene glycol molecules, 1 sulphate ion, 1 acetate molecule and 73 water molecules.

The atomic coordinates and structure factors have been deposited into the Brookhaven Protein Data Bank under the accession numbers 6S2W.

### S. pombe strains and growth media

The *S. pombe* strains used in this study are listed in Table S2. Strains were generated by transformation with a lithium acetate-based method. The *mmi1*Δ cells were generated from a parental strain possessing a deletion of *mei4+*, since the absence of Mmi1 leads to severe growth and viability defects due to the deleterious expression of Mei4. All experiments were performed using minimal medium (EMM Broth, Formedium, #PMD0210) supplemented with 150 mg/L of each adenine (Sigma, #A2786), L-histidine (Sigma, #H8000), uracil (Sigma, #U750) and L-lysine (Sigma, #L5501) but lacking L-leucine (EMM-LEU). To assess mating/sporulation efficiency, cells plated on EMM-LEU medium for 5 days at 30°C were exposed to iodine crystals (Sigma, #326143) for 5 min at room temperature.

### Co-immunoprecipitation and Total protein analyses

Experiments were performed as described in Simonetti et al, 2017 ^21^ except that detection was done with a Vilber Lourmat Fusion Fx7 imager.

### RNA extraction and RT-qPCR

Experiments were performed as described in Simonetti et al, 2017 ^21^ except that 100 units Maxima Reverse Transcriptase (Thermo Scientific, #EP0743) were used for reverse transcription reactions.

Oligonucleotides used in qPCR reactions are listed in Table S3.

### RNA-immunoprecipitation

Experiments were performed as described in Simonetti et al, 2017 ^21^ with the following modifications: 40 ODs of cells were grown to mid-log phase at 30°C in EMM-LEU and cross-linked with 0.2% formaldehyde for 20 min. Following quenching with 250 mM glycine for 5 min, cells were harvested by centrifugation. Cell pellets were resuspended in 2 ml RIPA buffer (50 mM Tris-HCl pH 8, 150 mM NaCl, 1% NP-40, 0.5% sodium deoxycholate, 0.1% SDS, 2 mM EDTA, 2 mM benzamidine, 1X Roche complete EDTA-free protease inhibitor cocktail and 80 U RNaseOUT Ribonuclease inhibitor (Invitrogen, #10777-019)) to make “pop-corn”. Lysis was performed using a Ball Mill (Retsch, MM400) for 15 min at 15 Hz frequency. Extracts were cleared by centrifugation before immunoprecipitation with 1 mg of pre-washed rabbit IgG-conjugated M-270 Epoxy Dynabeads (Invitrogen, #14311D) for 1 hour at 4°C. Beads were then washed once with low salt buffer (10 mM Tris-HCl pH7.5, 150 mM NaCl, 0.5% Triton X-100), twice with high salt buffer (10 mM Tris-HCl pH7.5, 1 M NaCl, 0.5% Triton X-100) and once again with low salt buffer for 10 min at room temperature. Total and immunoprecipitated RNAs were decrosslinked at 70°C for 45 min in the presence of reverse buffer (10 mM Tris-HCl pH 6.8, 5 mM EDTA, 10 mM DTT, 1% SDS) and treated with proteinase K for 30 min at 37°C. RNAs samples were next extracted with phenol:chloroform 5:1 pH4.7 (Sigma, #P1944), precipitated with ethanol and treated with DNase (Ambion, #AM1906) prior to RT-qPCR analyses.

### SmFISH

Quasar 670-labeled meiRNA probes were designed using Stellaris Probe Designer tool (Table S4) and synthesized by Biosearch Technologies. Single molecule RNA Fluorescence In-Situ Hybridization (smFISH) was performed according to the manufacturer’s protocol (Biosearch Technologies) with minor modifications.

Vegetative cells were plated on EMM-LEU and grown for 3 days at 30°C. Cells were then resuspended in 1X PBS containing 3.7% formaldehyde to an OD_600nm_ of 0.3, treated with Zymolyase 100 T for cell wall digestion and permeabilized in 70% ethanol prior to over-night incubation with meiRNA probes. Stellaris RNA FISH hybridization and wash buffers were obtained from Biosearch Technologies. DAPI stained cells were resuspended in Vectashield antifade mounting medium (Vector laboratories) and imaged using DM6000B Leica microscope with a 100X, numerical aperture 1.4 (HCX Plan-Apo) oil immersion objective and a charge-coupled device (CCD) camera (CoolSNAP HQ; Photometrics). Optical Z sections (0.2 µm step size, 25 sections) were acquired using a piezo-electric motor (LVDT; Physik Instrument) and the MetaMorph 6.1 software prior to maximum-intensity projection into a single plane. Images were processed in ImageJ (NIH).

## Acknowledgments

We acknowledge SOLEIL for provision of synchrotron radiation facilities and we would like to thank the beamline scientists for their assistance in using beamlines Proxima-1 and Proxima-2a. We are indebted to Dr Y. Mechulam for his help with data collection and to Pr. C. Gaillardin (INA-PG, Thiverval-Grignon, France) for the gift of the *S. pombe* cDNA library.

This work was supported by Ecole Polytechnique, the Institut de Biologie Intégrative de la Cellule, the Centre National pour la Recherche Scientifique and the Agence Nationale pour la Recherche [grants ANR-16-CE11-0003 to M.G and ANR-16-CE12-0031-01 to M.R.]. D.H. was supported by a PhD fellowship from the French Ministère de l’Enseignement Supérieur et de la Recherche (MESR).

## Authors’ Contributions

DH and VA performed experiments. DH, VA, BP, MR and MG designed the study, analyzed the data, drafted the article and approved the version to be published.

## Competing Interests

We have no competing interests.

