## Supplementary files for "Formation of *S. pombe* Erh1 homodimer mediates gametogenic gene silencing and meiosis progression"

**Table S1: Oligonucleotides and plasmids used to over-express proteins in *E. coli***

| ORF | Primer name | Primer sequence (the restriction sites are underlined) | Plasmid |
|---|---|---|---|
| Erh1 | oMG511 | TATAGGATCCAGCCCCCCACCCGCCG | pMG921 |
| | oMG512 | GCGGCTCGAGTTACGGAATCTGACGAGCCGC | |
| Erh1$_{I11R,L13R}$ | oMG629 | CGAATCTCATATCAGGCTGAGGATTCAGCAAGGTTCTGACCCT | pMG945 |
| | oMG630 | CCTTGCTGAATCCTCAGCCTGATATGAGATTCGGCGGGTGGGGGGC | |
| Mmi1-[95-122] | oMG605 | TATA GGATCC GGTAAATATGATTTTAGCAGGC | pMG915 |
| | oMG606 | TATA CTCGAG TCAAGACTCACGACGAAGG | |

**Table S2: *S. pombe* strains used in this study**

| Strain | Genotype |
|---|---|
| PR040 | h90, *ura4-DS/E, ade6-M210, leu1-32, mat3M::ura4+* |
| PR808 | PR040, *pREP41::LEU2* |
| PR1026 | PR040, mei4*::nat*$^{R}$*MX* mmi1*::hph*$^{R}$*MX kan*$^{R}$*MX*::P$_{nmt41}$-TAP-Mei2 pREP41*::LEU2* |
| PR1316 | PR040, mei4*::nat*$^{R}$*MX* mmi1*::hph*$^{R}$*MX* pREP41*::LEU2* |
| PR1413 | PR040, *kan*$^{R}$*MX*::P$_{nmt41}$-TAP-Mei2 pREP41*::LEU2* |
| PR1414 | PR040, erh1::*nat*$^{R}$*MX kan*$^{R}$*MX*::P$_{nmt41}$-TAP-Mei2 pREP41*::LEU2* |
| PR1415 | PR040, erh1::*nat*$^{R}$*MX kan*$^{R}$*MX*::P$_{nmt41}$-TAP-Mei2 pREP41-GFP-Erh1::*LEU2* |
| PR1416 | PR040, erh1::*nat*$^{R}$*MX kan*$^{R}$*MX*::P$_{nmt41}$-TAP-Mei2 pREP41-GFP-Erh1$_{I11R,L13R}$::*LEU2* |
| PR1420 | PR040, erh1::*nat*$^{R}$*MX* pREP41*::LEU2* |
| PR1421 | PR040, erh1::*nat*$^{R}$*MX* pREP41-GFP-Erh1::*LEU2* |
| PR1422 | PR040, erh1::*nat*$^{R}$*MX* pREP41-GFP-Erh1$_{I11R,L13R}$::*LEU2* |
| PR1440 | PR040, erh1::*nat*$^{R}$*MX* Mmi1-TAP::*hph*$^{R}$*MX* pREP41-GFP-Erh1::*LEU2* |
| PR1441 | PR040, erh1::*nat*$^{R}$*MX* Mmi1-TAP::*hph*$^{R}$*MX* pREP41-GFP- Erh1$_{I11R,L13R}$::*LEU2* |

**Table S3: Oligonucleotides used in this study**

| Primers | Sequence | Related figures |
|---|---|---|
| P249: *mei4*+ fwd | 5’-TGGATCAGATCCGTGGAATC-3’ | 2C, 2D |
| P250: *mei4*+ rev | 5’-AACGCTCGATTAGAAGGCAT-3’ | 2C, 2D |
| P253: *act1*+ fwd | 5’-AACCCTCAGCTTTGGGTCTT-3’ | 2C, 2D |
| P254: *act1*+ rev | 5’-TTTGCATACGATCGGCAATA-3’ | 2C, 2D |
| P325: *ssm4*+ fwd | 5’-ACACAGTTTACGGGATTCTA-3’ | 2C, 2D |
| P326: *ssm4*+ rev | 5’-GATTGTGATGAAAACTGGGT-3’ | 2C, 2D |

**Table S4: meiRNA probes for SmFISH**

| Probes | Sequence |
|---|---|
| #1 | 5’-ATACCCACTAAGTCTGTTTA-3’ |
| #2 | 5’-CGGCAGAAGATTGACCAACA-3’ |
| #3 | 5’-GCATATTCCGTCTTACAATA-3’ |
| #4 | 5’-ACCAACTAAAGCGATCTTGC-3’ |
| #5 | 5’-GACCATTTCAAAATGTTGCA-3’ |
| #6 | 5’-TACCGAATCCAGCTTTTTGA-3’ |
| #7 | 5’-CAGAGCTTAGAAGACAAGGT-3’ |
| #8 | 5’-TAACTGGACCCCATCAAGAA-3’ |
| #9 | 5’-TAAACCAACTTGGGGGTTGG-3’ |
| #10 | 5’-TCTAAGCTACTATTCATCCA-3’ |
| #11 | 5’-AGTAGATTCCATCAGTCATA-3’ |
| #12 | 5’-TGCAGCCAAAAAGTGTACCA-3’ |
| #13 | 5’-CATTGTAAGTGCTTTCAAGG-3’ |
| #14 | 5’-TTCAGTCATTCGCAAAGTTT-3’ |
| #15 | 5’-AGTCGTTTTATTTCTTTTCT-3’ |
| #16 | 5’-GTTTCAACAATAGTTCAGGT-3’ |
| #17 | 5’-TCTGTTTCAGGAATACGTTT-3’ |
| #18 | 5’-TGTTTCGCATCAAACTTTCA-3’ |
| #19 | 5’-GCGTTTAAACAAACTGCGGG-3’ |
| #20 | 5’-TGGTTTCAGCACGTTTTCAA-3’ |
| #21 | 5’-TTGGTTTGCAGGGTTTAACG-3’ |
| #22 | 5’-CTTGCTGTGGTTATTGTTTA-3’ |

***Expanded view Figure 1: Structural rearrangements of Erh1 upon binding to Mmi1***

A. Superimposition of the three copies of Erh1 present in the asymmetric unit.

B. Comparison of apo and Mmi1-bound Erh1 structures (rmsd values raging from 0.8-1.2Å over 80 Cα atoms). This reveals a large conformational change of the α2-β3 hinge characterized by a 4Å translation of Ile66 Cα atom and a 180° rotation of Tyr67 side chain (indicated by a red arrow). As a result, this renders accessible a hydrophobic cavity at the surface of Erh1, in which Phe99 side chain from one Mmi1 accommodates in the Erh1-bound structure. Concomitantly, Erh1 Tyr67 hydroxyl group forms an hydrogen bond (depicted as a black dashed line) with the Gly107 carbonyl group from the second Mmi1 peptide.

C. Upon Mmi1 binding, the N-terminal extremity of Erh1 strand β1 rearranges and His9 side chain flips by 180° (red arrow) to stack with Mmi1 Trp112 side chain.

***Expanded view Figure 2: Erh1 does not contribute to Mmi1-dependent downregulation of Mei2***

Western blot showing total TAP-Mei2 levels expressed from the $P_{nmt41}$ promoter in strains of the indicated genotypes. anti-GFP and anti-CDC2 antibodies were used to evaluate Erh1 levels and loading, respectively.

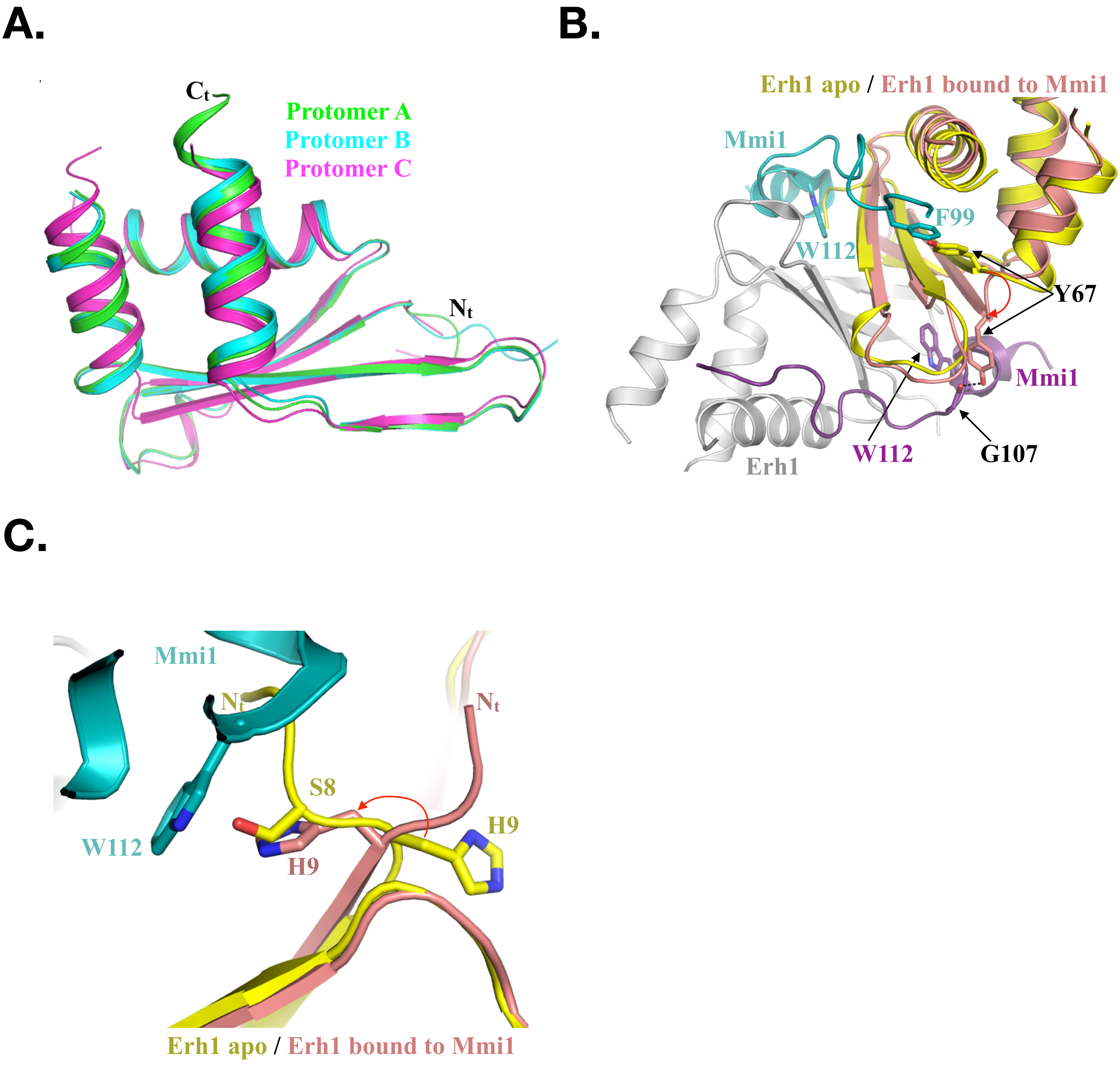


**Supplemental Figure 1: Structural rearrangements of Erh1 upon binding to Mmi1**

A.

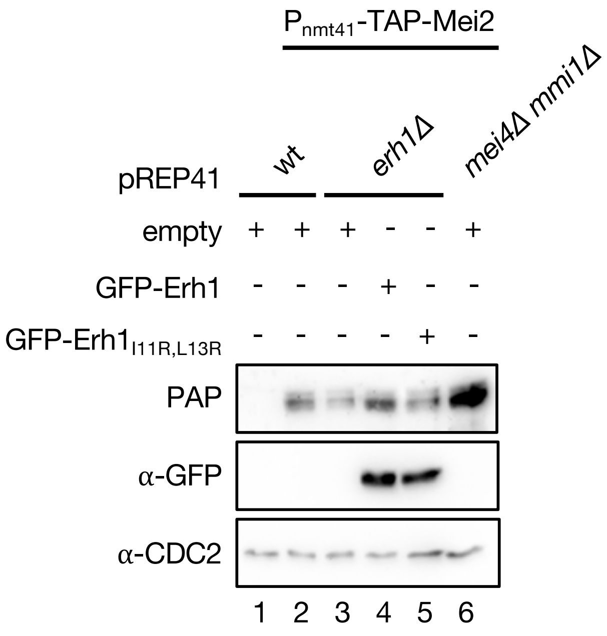


**Supplemental Figure 2: Erh1 does not contribute to Mmi1-dependent downregulation of Mei2**
